# Functional dispersion of wetland birds, invertebrates and plants more strongly influenced by hydroperiod than each other

**DOI:** 10.1101/2020.07.29.226910

**Authors:** Jody Daniel, Rebecca C Rooney

**Affiliations:** B2-251, Department of Biology, University of Waterloo, Waterloo, Ontario, Canada, N2L 3G1

**Keywords:** Structural equation model, hydroperiod, Prairie Pothole Region, biological communities, wetland, marsh, functional dispersion, permanence class

## Abstract

The relative role of biological and abiotic filters on the assembly of co-occurring taxa is widely debated. While some authors point to biological interactions (e.g., competition) as the stronger driver of ecological selection, others assert that abiotic conditions are more important because they filter species at the regional level. Because communities influenced by a dominant abiotic filter, (e.g., Prairie Pothole Region (PPR) wetlands, each varying in ponded water permanence), often have strong cross-taxon relationships, we can study these communities to better understand the relative influence of abiotic vs biotic filters on community structure. Using functional dispersion as our measure of communities, we test six alternate hypotheses about the relative importance of various pathways representing influence of biological and permanence filters on birds, aquatic macroinvertebrates and wetland plants in the northwest PPR using structural equation modeling. We aimed to understand whether: 1) ponded water permanence alone explained functional dispersion; 2) the influence of permanence on functional dispersion was direct or mediated; and 3) abiotic filtering by permanence was stronger than biotic filtering by co-occurring taxa. The best model suggests that there is a direct influence of permanence on the functional dispersion of each taxonomic group and that both bird and macroinvertebrate functional dispersion are causally related to plant functional dispersion, though for invertebrates the influence of plants is much less than that of permanence. Thus, the relative importance of wetland permanence and the functional dispersion of co-occurring taxa depends on which taxon is considered in PPR wetlands.

## Introduction

Though there is consensus among community ecologists that abiotic and biotic filters structure communities (Azeria et al. 2009, Qian and Kissling 2010, Chase and Myers 2011, Cabra-García et al. 2012, Devercelli et al. 2016), the relative role of each filter is widely debated (Kraft et al. 2015, Duan et al. 2016). Poff (1997) presented a nested filter conceptual model of community assembly. Poff argued that species within the regional species pool must first pass through the coarse filter of abiotic conditions; species with functional traits adapted to the range of conditions set by the abiotic filter would survive. Next, these surviving species would influence each other’s abundances through biological interactions – a more fine-scaled filter. Since Poff (1997), several other authors have found support for this nested filter model (e.g., Ackerly and Cornwell 2007, Williams et al. 2009, Aronson et al. 2016). Though manipulative experiments should prove useful in understanding the relative role of abiotic and biotic filters (Tiunov and Scheu 2005, Wardle 2006, Maynard et al. 2018), we posit that studying taxonomic groups exposed to a predominant environmental filter could help in partitioning their relative role in the assembly of communities by providing a simplified but environmentally relevant system.

Wetlands of varying ponded water permanence in the northwestern Prairie Pothole Region (PPR) provide an excellent candidate for such a model system. Wetlands in the PPR differ in the length of time ponded water is present (i.e., hydroperiod), some containing ponded water year-round and others drying up a few weeks after spring snowmelt (Stewart and Kantrud 1971, Leibowitz and Vining 2003). While the diversity and community structure of birds, aquatic macroinvertebrates and plants in PPR wetlands appear directly impacted by hydroperiod (Casanova and Brock 2000, Ruhí et al. 2014, Gleason and Rooney 2018, Daniel et al. 2019), interactions are evident among these taxa: 1) birds forage, roost in and nest on plants, but also disperse their seeds (e.g., Fox et al. 2011, Ayers et al. 2015, Soons et al. 2016); 2) birds consume aquatic macroinvertebrates and can influence their egg bank (e.g., Horváth et al. 2012; van Leeuwen et al. 2017); and 4) plants provide habitat for aquatic macroinvertebrates (e.g., Gleason et al. 2018). Evaluating the relative role of these filters can be pursued using a causal framework, which requires a univariate proxy of community composition; functional dispersion – a measure of how species abundances vary in trait space (Schleuter et al. 2010) – is a reliable proxy for community structure in studying community assembly processes (Gerhold et al. 2015). Dispersion is a preferred univariate measure of composition when studying these assembly processes because it captures how abiotic and biotic filtering can influence community structure. Species with similar functional traits will “pass through” an abiotic filter, resulting in low dispersion. In contrast, interspecific competition will encourage higher functional dispersion as species with different functional traits enable niche partitioning (Gerhold et al. 2015). We would expect, therefore, that if there is support for Poff’s nested filter model, then the influence of hydroperiod on a taxon’s functional dispersion would be stronger than the correlation in functional dispersion between taxa.

We can conceive of six distinct, plausible models to describe the possible interactions in our model system. First, the functional dispersion of taxanomic groups in our wetlands may be entirely the result of the influence of ponded water permanence on each taxon, independently (Fig. 1A). For example, hydroperiod may determine which plants in the seedbank will germinate and prolonged flooding can exclude ill adapted species (van der Valk 1981, Casanova and Brock 2000, Euliss et al. 2004, Tsai et al. 2012, Mushet et al. 2018). Hydroperiod may also filter macroinvertebrates lacking desiccation adaptations from lower permanence wetlands (Hall et al. 2004, Gleason and Rooney 2018). Birds also show sensitivity to hydroperiod; it can dictate which terrestrial birds, shorebirds and waterfowl can establish (Niemuth et al. 2006, Morissette et al. 2013), with waterbirds particularly responsive to the extent of open water (O’Neal et al. 2008). In this case, the functional dispersion of the three taxa are independent.

**Figure 1.**
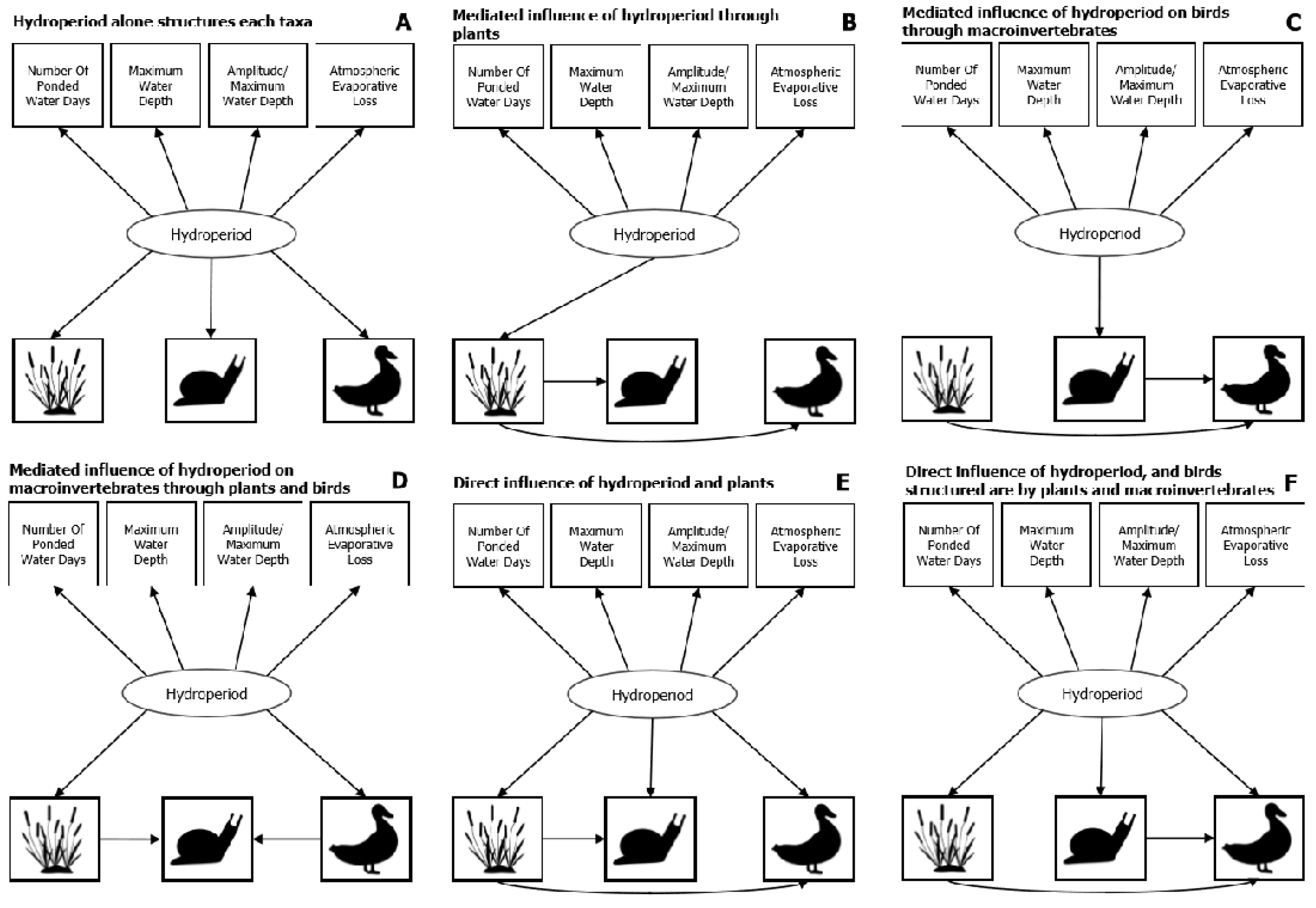
Six candidate structural equation model to evaluate the relative influence of biological interactions and hydroperiod on community congruence.

**Figure 2.**
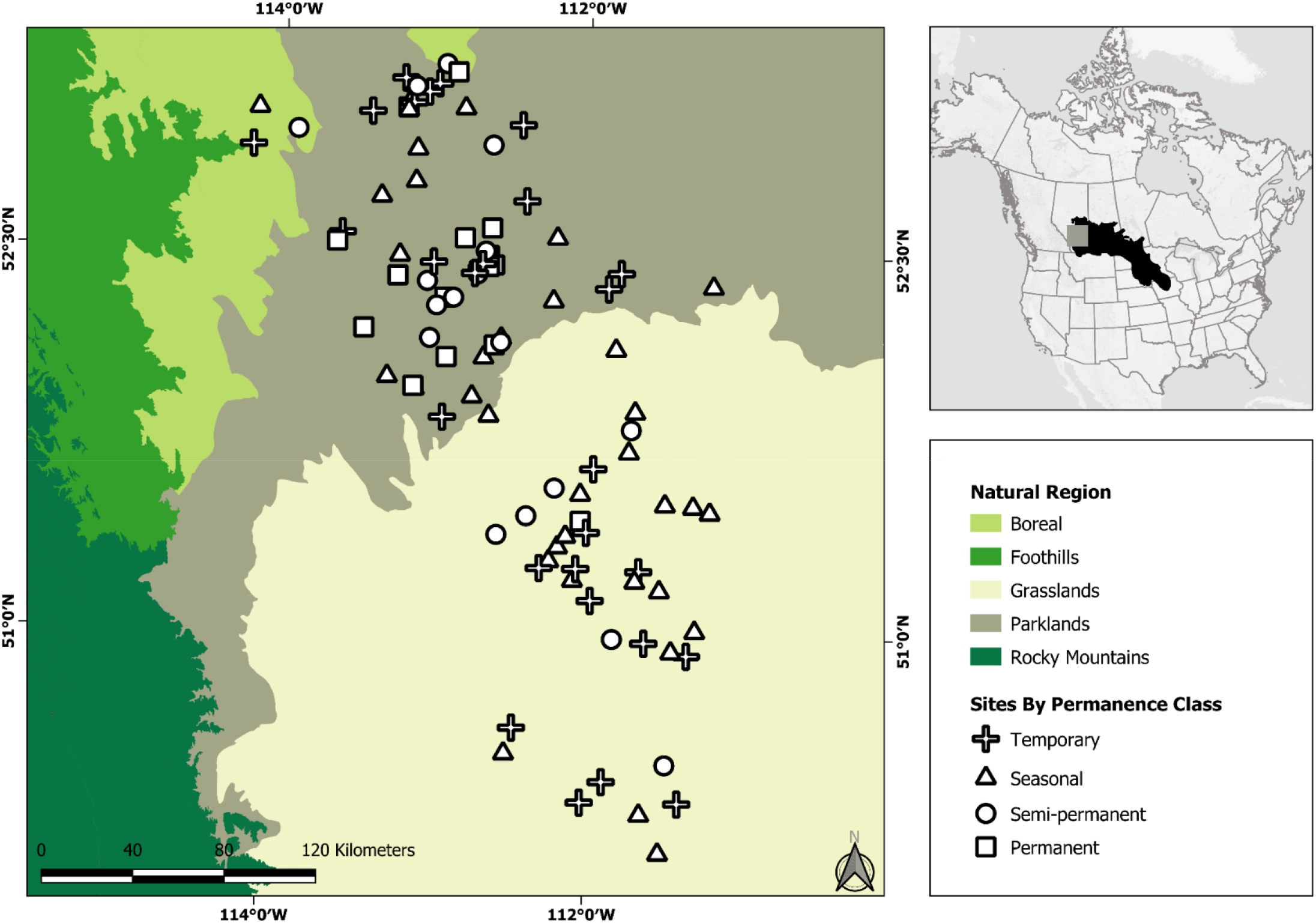
Map of our study sites and region, which are situated in the northern prairie pothole region (inset map). Our 96 wetland sites represented temporary (n = 28), seasonal (n = 35), semi-permanent (n = 17), and permanent (n = 14) ponded-water permanence classes, and they covered the Grassland and Parkland Natural Regions. The number of wetlands in each permanence class matched the frequency distribution of permanence classes in the central and southern wetland inventories (Government of Alberta 2014).

Alternatively, one or more taxon may be unaffected by hydroperiod directly, but its functional dispersion may be subject to indirect effects of ponded water permanence because it is directly influenced by the functional dispersion of a co-occurring taxon that itself is influenced by hydroperiod. We can conceive of three possible pathways that could yield this type of mediated effect. Our first mediated-effects model posits that hydroperiod structures the functional dispersion of wetland communities through its effect on the functional dispersion of plants (Fig. 1B). Here, we hypothesized that because plants determine whether birds can nest or forage (Klaassen and Nolet 2007, Austin and Buhl 2011, Fox et al. 2011, Ayers et al. 2015) and act as substrate for numerous aquatic macroinvertebrate families (Campeau et al. 1994, Lee Foote and Rice Hornung 2005), they shape resource availability for the other taxa. In this model, hydroperiod directly influences the functional dispersion of plants, as the moisture gradient is known to be a strong constraint on the distribution of hydrophytes (e.g., van der Valk 1981, Euliss et al. 2004), but only indirectly influences birds and macroinvertebrates.

For our second mediated-effects model (Fig. 1C), we posit that plants are independent of hydroperiod, and are rather structured by stochastic factors. Wetland plants are largely wind dispersed (Guarino et al. 2005) and plant communities are typically recruitment limited (e.g., Hurtt and Pacala 1995), hence the abundance of plants may prove random with respect to ecological selection pressures like hydroperiod. However, hydroperiod should directly structure macroinvertebrate functional dispersion, with less permanently ponded wetlands filtering out taxa that lack desiccation tolerance strategies (e.g., Gleason and Rooney 2018). In this model, bird functional dispersion is directly influenced by the functional dispersion of both plants and macroinvertebrates, since they structure the availability of roosting or nesting habitat and foraging opportunities for birds (e.g., Gurney et al. 2017, Vanausdall and Dinsmore 2019). More, because birds are highly mobile, they may prove the most sensitive to cross-taxon influences as they select wetlands for foraging and nesting based on the availability of preferred habitat and forage. Birds are thus indirectly influenced by hydroperiod, through its influence on aquatic macroinvertebrates.

Our final mediated model posits that hydroperiod structures both bird and plant functional dispersion, which in turn structure the functional dispersion of aquatic macroinvertebrates (Fig. 1D). For example, a study from Hungary reported that wetlands populated by waterbirds typically support smaller species of aquatic macroinvertebrates, which are less easily consumed by birds (Horváth et al. 2012). Similarly, vegetation, by providing habitat heterogeneity and refugia, is reported to influence the diversity and abundance and diversity aquatic macroinvertebrates (Davis and Bidwell 2008, Meyer et al. 2015; Gleason et al. 2018). Thus, the influence of hydroperiod on aquatic macroinvertebrates could be indirect, mediated through its effect on the functional dispersion of birds and plants in the wetland.

Yet, if Poff’s nested filter model is correct, we should see both a strong, direct influence of hydroperiod on the functional dispersion of each taxon and a simultaneous but weaker cross-taxon influence of functional dispersion, at least between some taxon pairs. We conceived of two competing, but plausible models that combine direct hydroperiod effects with direct cross-taxon effects. Our first combined-influence model (Fig. 1E) is similar to Fig. 1B, but with the addition of a direct influence of hydroperiod on both aquatic macroinvertebrates and birds, as is depicted in Fig. 1A. Our final combined-influence model (Fig. 1F), incorporates the direct effect of hydroperiod on the functional dispersion of all taxa (like in Fig. 1A) with the cross-taxon influence of the functional dispersion of plants and aquatic macroinvertebrates on plants, birds (as visualized in Fig. 1C. posits that the functional dispersion of all communities is structured by hydroperiod, but it is bird functional dispersion that is structured by aquatic macroinvertebrates and plants (Fig. 1F). We believe that this model may explain community functional dispersion because birds are the most transient of these taxa, and they can select wetlands for foraging and nesting based on whether their preferred habitat is present.

We tested our six competing hypotheses regarding how the functional dispersion values of birds, plants, and aquatic macroinvertebrates relate to hydroperiod and the functional dispersion of co-occurring taxa. We asked 1) whether functional dispersion is driven by hydroperiod alone (Fig. 1A) or by both hydroperiod and the functional dispersion of co-occurring taxonomic groups (Fig. 1B-F), 2) whether the influence of hydroperiod on each taxonomic group was primarily direct or indirect; and 3) whether hydroperiod had a stronger influence on the functional dispersion of taxonomic groups than did the functional dispersion of co-occurring taxonomic groups (i.e., support for Poff’s nested filter model; Fig. 1E,F). We used structural equation modelling and AIC to evaluate the support for our six alternative hypotheses (Fig. 1). We predicted that there would be support for Poff’s nested filter model, which would be evidenced by 1) the influence of wetland hydroperiod exceeding that of the functional dispersion of co-occurring taxonomic groups, as this theory dictates that abiotic filters like hydroperiod should take precedence over biological filters and 2) a direct influence of both hydroperiod and cross-taxon interactions on community functional dispersion.

## Methods

### Study Area

Our wetlands are prairie potholes in the Grassland and Parkland Natural Regions of Alberta, Canada (Fig. 1). They are depressions that fill with ponded water, which were formed in the last glacial period (Wright 1972). Also in this region, the climate is semi-arid, as the rate of potential evapotranspiration exceeds that of annual precipitation (Hayashi et al. 2016). In the Grassland Natural Region, the dominant vegetation is mixed-grass prairie. Conversely, in the Parkland Natural Region, deciduous trees and grasses dominate (Downing and Pettapiece 2006).

### Study Design

We surveyed 96 wetlands that ranged in pond permanence class (sensu Stewart and Kantrud 1971) from temporary with a hydroperiod on the order of weeks to permanent with ponded water year round, even in dry years. We selected sites to mirror the frequency distribution of permanence classes in the Alberta Merged Wetland Inventory (Government of Alberta 2014), and so included an unequal number of wetlands per permeance class category. Generally, most of our wetlands were small (mean size 0.81 ± 0.12 SE ha), reflecting the dominance of small prairie pothole wetlands in the region, independent of their permanence class.

### Biological Surveys

#### Birds

We used visual and auditory point counts to survey birds twice during the peak breeding season (May-June in either 2014 or 2015) to record the presence of birds actively foraging or breeding (singing, nesting, territorial displays) in the study wetlands. Fly-overs were excluded from data analyses. In summary, visual surveys commenced first; they lasted for 10 minutes and were carried out from a vantage that allowed a clear view of the open water zone. We next conducted an 8-minute auditory survey. These surveys were 100-m, fixed-radius point counts, occurring at the center of the wetland. When a wetland was larger than 3 ha, we conducted multiple auditory surveys; each point-count location was at least 100 m from the wetland edge and 200 m from any other point-count location. We summed abundances across the multiple auditory point counts to account for differences in wetland size. For both visual and auditory counts, we recorded the identity and abundance of species (species list in Appendix S1A). Importantly, species abundances were summed across visits, rather than averaged, to account for the staggered breeding seasons among species. Additional details on the bird surveys are reported in Anderson and Rooney (2019).

#### Aquatic Macroinvertebrates

We applied a revised version (Gleason and Rooney 2017) of the quadrat-column-core method (Meyer et al. 2013) to survey aquatic macroinvertebrates. We sampled aquatic macroinvertebrates in both the open water (submersed and floating vegetation) and the emergent (cattail, bulrush, or other robust perennial sedges) zones, when both were present. In each zone, we collected three replicates of a: 1) vigorously washed and clipped 0.25 m^2^ vegetation sample, from the emergent or submersed aquatic vegetation; and 2) two, 10-cm diameter water column samples. We composited the replicates of each sample type; this yielded one water column, sediment core, and vegetation sample in each wetland vegetation zone (open water and emergent). Following, for water column samples, we sorted aquatic macroinvertebrates to identify them to the lowest practical taxonomic level (typically Family), following Clifford (1991) Merrit et al. (2008). We used a Marchant box to sub-sample the vegetation sample, which was based on the protocol of the Canadian Aquatic Biomonitoring Network (Environment Canada 2014). Here, taxon abundances are area-weighted to estimate density per meter-squared. For both vegetation and water column samples, aquatic invertebrates were scaled to the meter squared. Next, we summed densities to represent each wetland zone, and subsequently averaged across zones for wetland-level invertebrate relative abundances. Additional information on the aquatic macroinvertebrates sampling are reported in Gleason and Rooney (2017), and a comprehensive taxonomic list is provided in Appendix S1B.

#### Plants

We conducted plant surveys during peak aboveground biomass (late July to August). During peak biomass, the presence of inflorescences allows for herbaceous plants to be confidently identified and cover values are at their maximum. We first delineated the wetland boundary based on the 50:50 rule for vegetation classification. After mapping the extent of each plant assemblage based on their vegetative structure (e.g., deciduous tree, forb, floating-leaved vegetation), we then mapped them by which species were co-dominant or dominant. We used a GPS/GNSS unit (SX Blue II receiver, by Geneq Inc., Montreal, Canada) to map these assemblages. For communities sized between 100-5000 m^2^, we identified the percentage cover (modified Braun-Blanquette approach) of each vascular plant species within five, 1 m^2^ quadrats. When communities were larger than 5000 m^2^, we surveyed an additional quadrat per 1000 m^2^ of plant community area. We also recorded the percentage cover of algae, bryophytes, bare ground, litter, rock, seedling/unidentified forb, standing dead litter, and open water (species list in Appendix S1C), but these cover classes were not included in subsequent analyses of vascular plant cover. For more details on plant survey methods, see Bolding et al. (2020).

### Characterizing Prairie Pothole Permanence

Wetland permanence is a latent variable that cannot be directly measured but can be quantified nonetheless by a set of correlated indicators within a structural equation modelling framework (Grace et al. 2010). We used four proxies for wetland permanence, all related to the concept of hydroperiod. We used a matrix including the :1) approximate number of days a wetland contained ponded water based on ca. biweekly staff gauge measurements during the open water season (May-September), 2) the maximum water depth observed during the survey period 3) water amplitude (i.e., maximum – minimum observed water depths) standardized by maximum water depth observed during the survey period, and 4) an index of evaporative losses to the atmosphere relative to water inputs based on stable isotopes analysis. For details on the stable isotope analysis, see Meyers (2018).

### Statistical Analysis

#### Calculating Functional Dispersion

We used functional dispersion as a proxy for community structure in studying community assembly process. Generally, when functional dispersion is high, some functional traits are more abundant that others indicating low evenness among functional traits; if abundances of different functional traits within a community are equal, we will see low functional dispersion (Finke and Snyder 2008, Comte et al. 2016). To measure the functional dispersion of each taxon, we used Rao’s quadratic entropy with the dBF function in the FD package (Laliberte et al. 2014) in R (R Core Team 2019). This functional dispersion index uses the weighted mean distance of each species to the group centroid, where weights are based on species relative abundances. Thus, functional dispersion is the variance in a species’ traits and where they are located in trait space (Schleuter et al. 2010), and it uses both species relative abundances and the pairwise functional differences to summarize functional diversity. Importantly, this index is not influenced by richness (Rao 1982).

The traits we select for functional diversity indices can have profound effects on our understanding of a communities’ ecology (Zhu et al. 2017); this suggests that our trait selection can affect our interpretation of the relative influence of permanence and cross-taxon interactions on functional dispersions. Because we aimed to capture as many assembly processes as possible, and not bias our analyses to traits related to abiotic filtering (Spasojevic and Suding 2012), we selected a wide range of traits and not simply those likely to be sensitive to wetland hydroperiod. For birds, we selected functional traits indicative of feeding behavior (e.g., ground gleaner, dabbler) nesting ecology (e.g., bank, reed), primary habitat (e.g., shoreline, grassland), wetland status (e.g., obligates versus facultative) and migratory status (e.g., neo-tropical migrant) (Appendix S2A). With macroinvertebrates, we used traits on feeding (e.g. filter feeder), behavior class (e.g., climbers) and desiccation strategy (e.g., disperser) (Appendix S2B). For plants, however, we used wetland indicator status (e.g., emergent), dispersal (e.g. wind), reproduction (e.g. vegetative), and nativity (e.g., native vs exotic) (Appendix S2C).

#### Partitioning the Influences of Community Composition into Environmental and Biological Components

We used structural equation models to evaluate the pathways that could explain the relative influence of abiotic and biotic filters on functional dispersion. We compared the fit of six candidate models. Our first model examined whether functional dispersion was explained by permanence alone, our second to forth models assessed whether there was a direct or indirect influence of permanence and our fifth to sixth models were assessments of the relative influence of permanence and biological interactions.

We implemented the structural equation models in the lavaan package (Rosseel 2012) of R (R Core Team 2019). Before implementing each model, we relativized each variable by their respective maximum values because they differed in range and scale. We were confident that our relativized data did not violate the assumption of multivariate normality, which is required for structural equation models, based on results of a Mardia’s multivariate kurtosis of multiple variables test (z = −0.325, *p*-value = 0.774), implemented using the mardiaKurtosis function in the semTools package (Jorgensen et al. 2018) in R (R Core Team 2019). For each model, we set the endogenous covariances to zero, fixed the factor loading of the approximate number of days that a wetland contained ponded water to 1.0 and used an unbiased estimator (wishart) for maximum likelihood estimation. To rank the candidate models, we used AIC_c_ and model fit statistics. Finally, we standardized all parameter estimates, to ensure that we could compare the relative influence of hydroperiod and biological interactions on each taxonomic group’s functional dispersion.

## Results

### Partitioning the Influences of Community Composition into Environmental and Biological Components

We compared the fit of six structural equation models (Fig. 1), each representing a different hypothesis about the relative influence of co-occurring taxonomic groups and ponded-water permanence on the functional dispersion of birds, aquatic macroinvertebrates and vascular plants in our study wetlands.

Our best model was model 1E, which hypothesized a direct influence of permanence on the functional dispersion of all three taxa and a further influence of plant functional dispersion on the functional dispersion of birds and aquatic macroinvertebrates (Table 1). Because no models were within two ΔAIC units of this top-performing model (Arnold 2010), we conclude that there is strong support for this model. More, the AIC weights indicate that a direct influence of permanence and plant functional dispersion on the functional dispersion of birds and aquatic macroinvertebrates was substantially more likely (AIC weight = 98), given the data, than the other five models (Table 1; Wagenmakers and Farrell 2004). Based on the p-value and chi-square statistic for this model (Table 1), we are confident there is strong support for this model, and it fit our data well. The standardized regression coefficients for this model are presented in Fig. 3.

**Table 1 –.** Performance of six candidate structural equation models predicting the relative influence of biological interactions and hydroperiod on community congruence. Direct influence of permanence and plants was the best model with the ΔIC of all other models being > 9 units.

| Model | Chi-square | *p*-value | AIC | $\Delta AIC$ | AIC weight |
| --- | --- | --- | --- | --- | --- |
| A: Hydroperiod alone structures taxon | 30.983 | 0.006 | -231.165 | 14.291 | 0.077 |
| B: Mediated influence of hydroperiod through plants | 42.022 | 0.000 | -219.990 | 25.465 | 0.000 |
| C: Mediated influence of hydroperiod through inverts on birds | 24.093 | 0.020 | -234.134 | 11.321 | 0.341 |
| D: Mediated influence of hydroperiod on birds through plants and inverts | 24.745 | 0.025 | -235.475 | 9.980 | 0.667 |
| E: Direct influence of hydroperiod and plants | 12.902 | 0.376 | -245.455 | 0.000 | 98.034 |
| F: Direct influence of hydroperiod, and birds are structured by plants and inverts | 22.220 | 0.035 | -236.030 | 9.426 | 0.880 |

**Figure 3.**
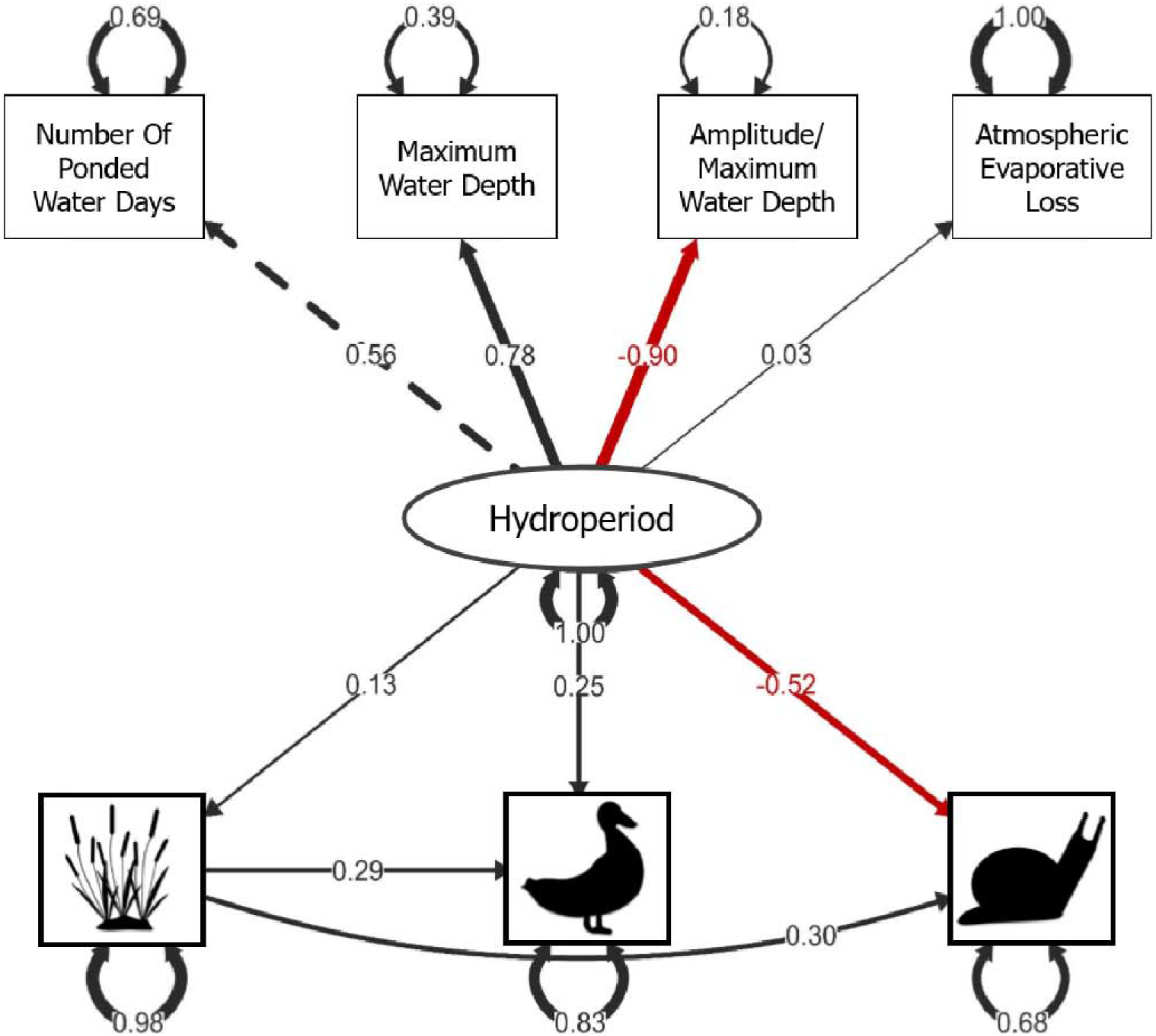
Measurement model for the best fitting of the six candidate structural models (there is a direct influence of permanence on the functional dispersion of birds, aquatic macroinvertebrates and plants, and plants directly structure birds and aquatic macroinvertebrates functional dispersion). Unidirectional lines are standardized regression slopes while bi-directional lines are standardized variances. The dashed, unidirectional line indicates which exogenous variable was fixed to a factor loading of one. Lines in black indicate that there was a positive relationship, while red lines indicate that the relationship was negative. Line thickness reflects the magnitude of the standardized regression slopes.

While permanence alone did not explain functional dispersion of our three wetland taxa (research question one), there was a direct influence of permanence on the functional dispersion of each taxon (research question two). Interestingly, the relative influence of permanence and the functional dispersion of plants on the functional dispersion of birds and invertebrates differed, depending on the taxon considered (research question three). Plant functional dispersion was solely influenced by hydroperiod. Hydroperiod had a much stronger influence on aquatic macroinvertebrate functional dispersion (standardized regression coefficient = −0.52) than did plant functional dispersion (standardized regression coefficient = 0.30). In contrast, for birds, the influence of hydroperiod (standardized regression coefficient = 0.25) was slightly weaker than that of plant functional dispersion standardized regression coefficient = (0.29), even when incorporating its indirect effect through plants (direct + indirect relative influence = 0.288). Generally, we conclude that the abiotic filter of hydroperiod is equal to or greater an influence on functional dispersion in each of our wetland taxa than the functional dispersion of co-occurring taxa, but the results are contingent on the taxon of interest.

## Discussion

Using the functional dispersion of co-occurring taxonomic groups in PPR wetlands, we evaluated whether there was support for Poff’s nested filter model in describing community assembly processes. We assumed that support for Poff’s model would be evidenced by a stronger direct influence of hydroperiod on functional dispersion than cross-taxon interactions. We find limited support for Poff’s nested filter model – abiotic filtering primarily influences functional dispersion and biotic filters are of lesser importance, at least for plants and aquatic macroinvertebrates. In birds, the functional dispersion of plants and the hydroperiod exerted similar influence. We asked three questions: 1) does the abiotic filter of ponded-water permanence alone structure prairie pothole wetland communities, 2) is the influence of this abiotic filter primarily direct or indirect and mediated through biological interactions, 3) what is the relative influence of this abiotic filter and the biological filtering of co-occurring communities?

To address our first question, we report that both the functional dispersion of co-occurring taxa and hydroperiod are important determinants of functional dispersion in these wetlands. This conclusion is supported by both observational studies and manipulative experiments, which report that biological interactions and environmental conditions structure resource availability for establishing taxa (e.g., Tiunov and Scheu 2005, Wardle 2006, Maynard et al. 2018). Environmental conditions can influence resource availability for primary producers by limiting whether nutrients necessary for growth are present (Fourqurean et al. 1992, Bowman et al. 1993, Guignard et al. 2017); and for consumers, environmental conditions determine whether energy gained (i.e., from feeding) (e.g., Schoo et al. 2012) is lower than the energetic costs to establish (e.g., from maintaining optimal body temperature, or time and effort placed into foraging) (Magnuson et al. 1979, Reid and Sprules 2018). Biological interactions, however, can influence resource availability through prey availability (for consumers) (Spivak et al. 2009, Groendahl and Fink 2017) or habitat provisioning (Thompson et al. 1996, Jackson et al. 2008). Thus, in predicting environmental change (Urban et al. 2016) or even species ranges (Dormann et al. 2018, Palacio and Girini 2018), we must also consider biological interactions.

To address our second question, whether the influence of hydroperiod was mainly direct or was mediated through the functional dispersion of co-occurring taxa, we find that the influence of hydroperiod was primarily direct. The functional dispersion of plants was driven exclusively by hydroperiod (standardized regression coefficient = 0.13). The functional dispersion of birds and aquatic macroinvertebrates included both a direct influence of hydroperiod (standardized regression coefficient = 0.25 and −0.52, respectively) and a much smaller indirect component. This mediated effect was through the influence of plant functional dispersion on birds (indirect standardized regression coefficient = 0.0377) and on aquatic macroinvertebrates (indirect standardized regression coefficient = 0.0390). Consequently, we conclude that taxon community structure in prairie pothole wetlands is mainly influenced by hydroperiod directly, even where indirect pathways of influence are supported by the data.

In terms of our third question, if there was support for Poff’s nested filter model of community assembly, wherein abiotic filters take primacy over secondary biological filters, we find moderate support for Poff. Hydroperiod was the most influential factor in determining the functional dispersion of our wetland taxa, but the functional dispersion of plants served to exert a secondary influence. In the case of birds, however, the influence of plant functional dispersion was slightly larger than the direct effect of wetland permanence, and on par with its combined direct and indirect effect. Hydroperiod appears to first filter out species from the regional species pool that lack the necessary adaptations or tolerances to persist, and only subsequently do biological interactions constrain community assembly. We can conclude that biotic filters may be less influential than abiotic filters in community assembly processes.

Though our model fit our data well, we could not account for some community assembly processes. Unknown is whether our hypothesized direct effects are not, in fact, mediated effects that we have categorized as direct effects. For instance, wetlands with longer hydroperiods may come to have higher bird functional dispersion because they are larger and some microhabitats are more abundant (Kantrud and Stewart 1984). If microhabitat availability is the direct pathway influence bird functional dispersion, the influence of hydroperiod would be best described as indirect, contrary to our findings. An additional missing link in our model is intra-taxon interactions. Widely debated before Poff’s (1997) nested filter model was introduced is Diamond’s (1975) assertion that the “checkered” distribution of species could be explained by competition-driven assembly. Though Diamond’s (1975) hypothesis sparked debates lasting several decades (e.g., Connor and Simberloff 1979, Gotelli 2000), some authors have found support for this model (e.g., Gotelli and McCabe 2002, Gotelli and Rohde 2002, Maestre et al. 2008). Because our models focused on cross-taxon interactions, we were unable to incorporate the influence of intraspecific competition as a community assembly process, which should be included in future studies.

Our model also excludes other possible drivers of community assembly [(e.g., influence of proximity to other wetlands on the regional species pool (Lokemoen and Woodward 1992, Galatowitsch 2006); water chemistry as an abiotic filter on aquatic macroinvertebrates (Longcore et al. 2006, Maurer et al. 2014); and influence of physio-chemical conditions on plants (Roy et al. 2019, Kraft et al. 2019)]. Inclusion of such factors could improve the predictive power regarding functional dispersion, though based on prior research (e.g.,, Gleason and Rooney 2017, Kraft et al. 2019, Anderson and Rooney 2019) we predict that hydroperiod will remain the most important environmental filter in our study system. Future modeling should investigate the relative importance of these factors in parsimoniously describing community assembly in PPR wetlands.

### Conclusion

Using structural equation modelling, we demonstrate that the functional dispersion of birds, aquatic macroinvertebrates and plants are explained by both hydroperiod and the functional dispersion of co-occurring taxa. Because environmental conditions generally had a stronger influence, even if indirect effects are considered, we find support for Poff’s nested filter model in the community assembly of co-occurring birds, vegetation and aquatic macroinvertebrates in PPR wetlands.

## Acknowledgements

We are grateful to Alberta Innovates grant #2094A and the Ontario Trillium Scholarship for funding this research. We are also thankful to Drs. Michael Anteau, Roland Hall and Derek Robinson for their feedback on an earlier draft of this manuscript and to two anonymous reviewers for their feedback. We extend thanks to the numerous field assistants on this project: Daina Anderson, Brandon Baer, Matt Bolding, Graham Howell, Adam Kraft, Jennifer Gleason, and Nicole Meyers and Heather Polan. To Dr. Erin Bayne, who supplied the automated recording units used to verify auditory surveys, we also extend thanks.

**Appendix S1A.** Code, scientific name, and common name of bird taxa observed at the 96 wetland sites.

| **Name** | **Scientific Name** | **Common Name** |
| --- | --- | --- |
| ALFL | *Recurvirostra americana* | Alder Flycatcher |
| AMAV | *Recurvirostra americana* | American Avocet |
| AMBI | *Botaurus lentiginosus* | American Bittern |
| AMCO | *Fulica americana* | American Coot |
| AMCR | *Corvus brachyrhynchos* | American Crow |
| AMGO | *Spinus tristis* | American Goldfinch |
| AMRE | *Setophaga ruticilla* | American Redstart |
| AMRO | *Turdus migratorius* | American Robin |
| AMWI | *Anas americana* | American Wigeon |
| BAIS | *Ammodramus bairdii* | Baird's Sparrow |
| BAOR | *Icterus galbula* | Baltimore Oriole |
| BARS | *Hirundo rustica* | Barn Swallow |
| BBMA | *Pica hudsonia* | Black-billed Magpie |
| BCCH | *Poecile atricapillus* | Black-capped Chickadee |
| BHCO | *Molothrus ater* | Brown-headed Cowbird |
| BLJA | *Cyanocitta cristata* | Blue Jay |
| BLTE | *Chlidonias niger* | Black Tern |
| BNST | *Himantopus mexicanus* | Black-necked Stilt |
| BOGU | *Chroicocephalus philadelphia* | Bonaparte's Gull |
| BRBL | *Euphagus cyanocephalus* | Brewer's Blackbird |
| BRTH | *Toxostoma rufum* | Brown Thrasher |
| BUFF | *Bucephala albeola* | Bufflehead |
| BWTE | *Anas discors* | Blue-winged Teal |
| CANG | *Branta canadensis* | Canada Goose |
| CANV | *Aythya valisineria* | Canvasback |
| CCSP | *Spizella pallida* | Clay-colored Sparrow |
| CEDW | *Bombycilla cedrorum* | Cedar Waxwing |
| CHSP | *Spizella passerina* | Chipping Sparrow |
| CITE | *Anas cyanoptera* | Cinnamon Teal |
| COGR | *Quiscalus quiscula* | Common Grackle |
| COLO | *Gavia immer* | Common Loon |
| CORA | *Corvus corax* | Common Raven |
| COYE | *Geothlypis trichas* | Common Yellowthroat |
| DOWO | *Picoides pubescens* | Downy Woodpecker |
| EAGR | *Podiceps nigricollis* | Eared Grebe |
| EAKI | *Tyrannus tyrannus* | Eastern Kingbird |
| EAPH | *Sayornis phoebe* | Eastern Phoebe |
| EUST | *Sturnus vulgaris* | European Starling |
| FISP | *Spizella pusilla* | Field Sparrow |
| FRGU | *Leucophaeus pipixcan* | Franklin's Gull |
| GADW | *Anas strepera* | Gadwall |
| GBHE | *Ardea herodias* | Great Blue Heron |
| GRCA | *Dumetella carolinensis* | Gray Catbird |
| GRSP | *Ammodramus savannarum* | Grasshopper Sparrow |
| GWTE | *Anas crecca* | Green-winged Teal |
| HAWO | *Picoides villosus* | Hairy Woodpecker |
| HOGR | *Podiceps auritus* | Horned Grebe |
| HOLA | *Eremophila alpestris* | Horned Lark |
| HOSP | *Passer domesticus* | House Sparrow |
| HOWR | *Troglodytes aedon* | House Wren |
| KILL | *Charadrius vociferus* | Killdeer |
| LBCU | *Numenius americanus* | Long-billed Curlew |
| LCSP | *Ammodramus leconteii* | Le Conte's Sparrow |
| LEFL | *Empidonax minimus* | Least Flycatcher |
| LESA | *Calidris minutilla* | Least Sandpiper |
| LESC | *Aythya affinis* | Lesser Scaup |
| LEYE | *Tringa flavipes* | Lesser Yellowlegs |
| LISP | *Melospiza lincolnii* | Lincoln's Sparrow |
| MAGO | *Limosa fedoa* | Marbled Godwit |
| MALL | *Anas platyrhynchos* | Mallard |
| MERL | *Falco columbarius* | Merlin |
| MODO | *Zenaidura macroura* | Mourning Dove |
| NESP | *Ammodramus nelsoni* | Nelson's Sparrow |
| NOFL | *Colaptes auratus* | Northern Flicker |
| NOHA | *Circus cyaneus* | Northern Harrier |
| NOPI | *Anas acuta* | Northern Pintail |
| NRWS | *Stelgidopteryx serripennis* | Northern Rough-winged Swallow |
| NSHO | *Anas clypeata* | Northern Shoveler |
| OVEN | *Seiurus aurocapilla* | Ovenbird |
| PIWO | *Dryocopus pileatus* | Pileated Woodpecker |
| RBGR | *Pheucticus ludovicianus* | Rose-breasted Grosbeak |
| RBGU | *Larus delawarensis* | Ring-billed Gull |
| RBNU | *Sitta canadensis* | Red-breasted Nuthatch |
| RCKI | *Regulus calendula* | Ruby-crowned Kinglet |
| REDH | *Aythya americana* | Redhead |
| REVI | *Vireo olivaceus* | Red-eyed Vireo |
| RENH/RHNP | *Phalaropus lobatus* | Red-necked Phalarope |
| RNDU | *Aythya collaris* | Ring-necked Duck |
| RTHA | *Buteo jamaicensis* | Red-tailed Hawk |
| RUBL | *Euphagus carolinus* | Rusty Blackbird |
| RUDU | *Oxyura jamaicensis* | Ruddy Duck |
| RUGR | *Bonasa umbellus* | Ruffed Grouse |
| RWBL | *Agelaius phoeniceus* | Red-winged Blackbird |
| SACR | *Antigone canadensis* | Sandhill Crane |
| SAVS | *Passerculus sandwichensis* | Savannah Sparrow |
| SORA | *Porzana carolina* | Sora |
| SOSP | *Melospiza melodia* | Song Sparrow |
| SPPI | *Anthus spragueii* | Sprague's Pipit |
| SWHA | *Buteo swainsoni* | Swainson's Hawk |
| SWSP | *Melospiza georgiana* | Swamp Sparrow |
| SWTH | *Catharus ustulatus* | Swainson's Thrush |
| TEWA | *Oreothlypis peregrina* | Tennessee Warbler |
| TRES | *Tachycineta bicolor* | Tree Swallow |
| UNKN | Unknown | Unknown |
| UPSA | *Bartramia longicauda* | Upland Sandpiper |
| VESP | *Poocetes gramineus* | Vesper Sparrow |
| WAVI | *Vireo gilvus* | Warbling Vireo |
| WEME | *Sturnella neglecta* | Western Meadowlark |
| WILL | *Tringa semipalmata* | Willet |
| WIPH | *Phalaropus tricolor* | Wilson's Phalarope |
| WISN | *Gallinago delicata* | Wilson's Snipe |
| WTSP | *Zonotrichia albicollis* | White-throated Sparrow |
| YEWA | *Setophaga petechia* | Yellow Warbler |
| YHBL | *Xanthocephalus xanthocephalus* | Yellow-headed Blackbird |

**Appendix S1B.**
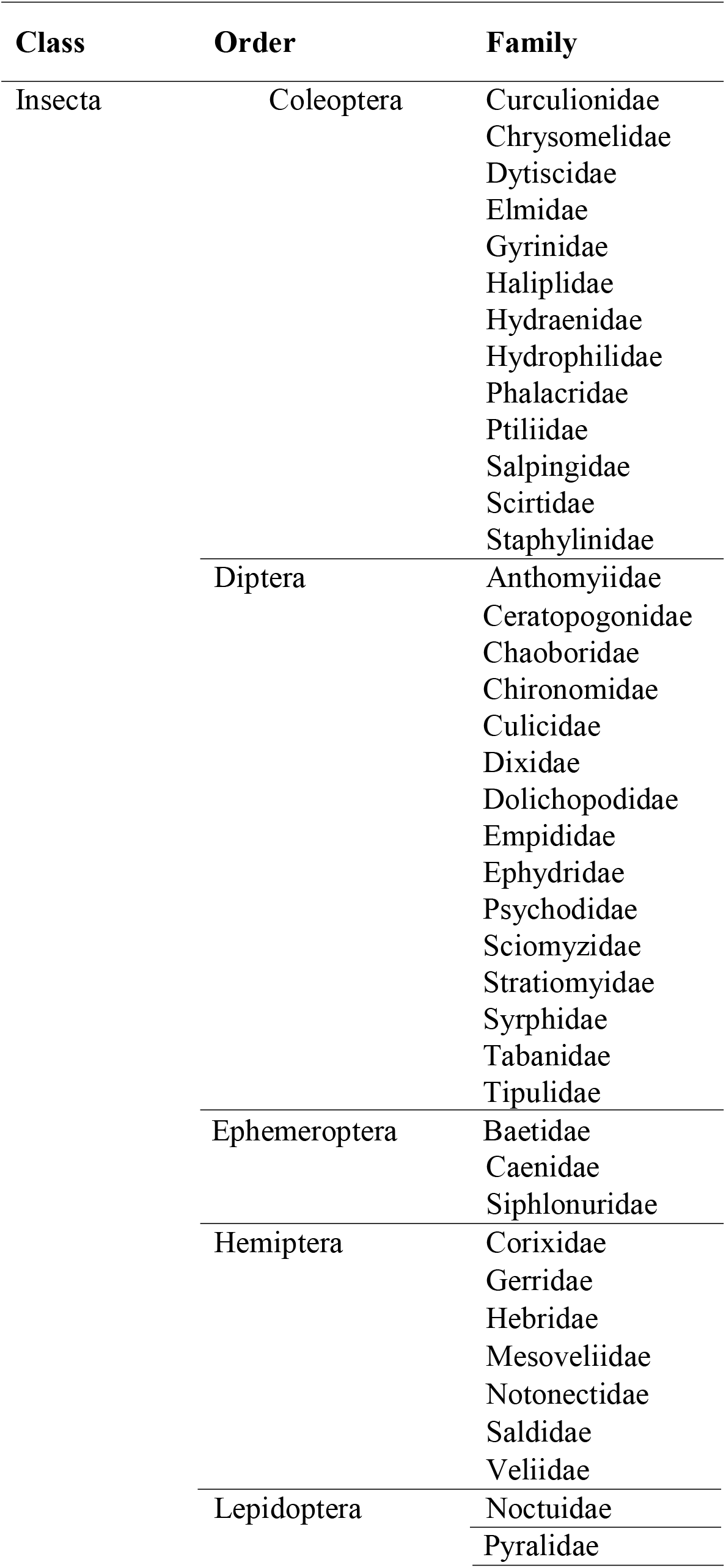
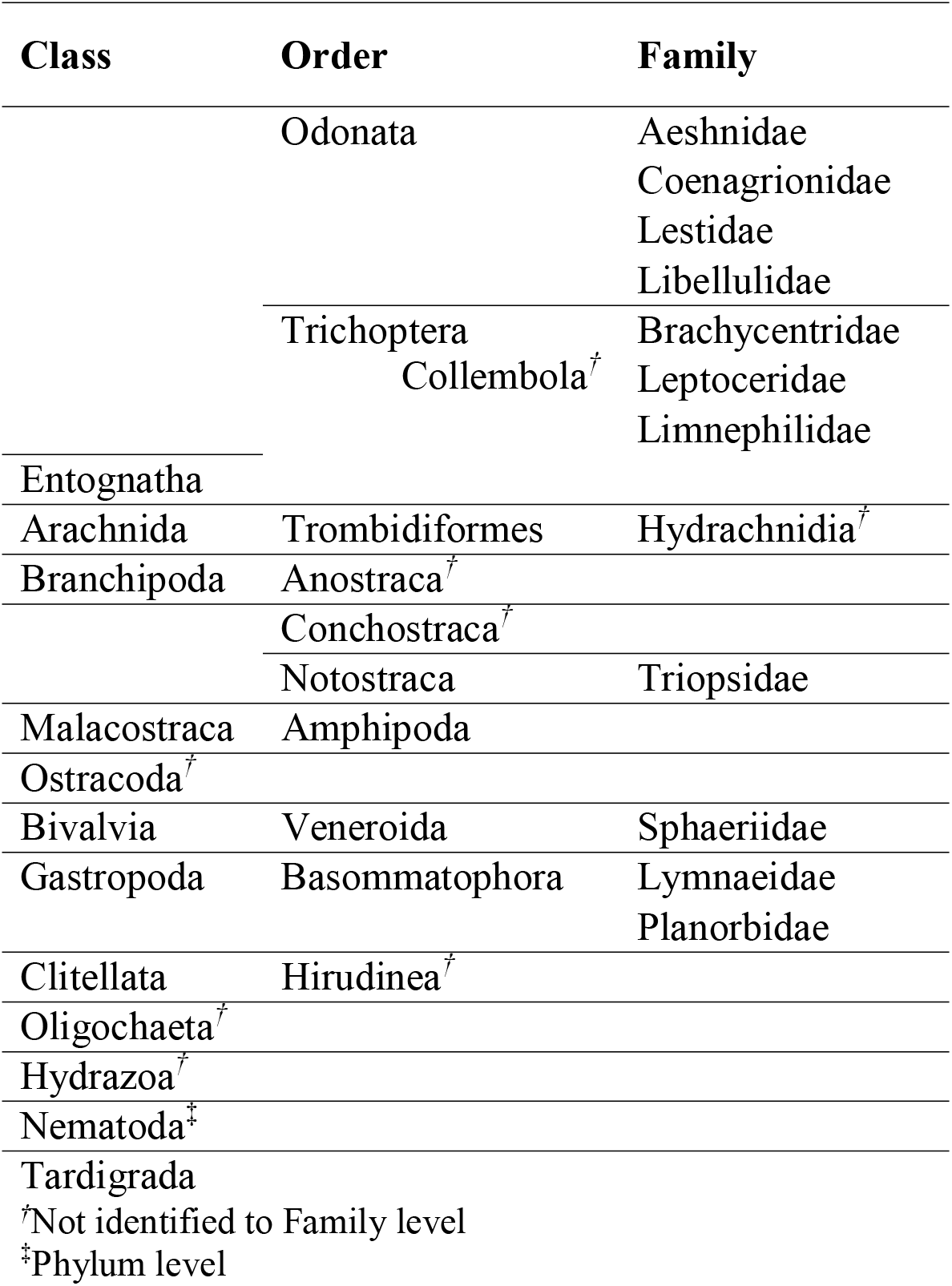
A list of all aquatic macroinvertebrate taxa identified with taxonomic resolution.

**Appendix S1C.** Code, scientific name, and common name of plant taxa observed at the 96 wetland sites.

| Code | Scientific Name | Common Name |
| --- | --- | --- |
| ACHALPIN | *Achillea alpina* | Siberian Yarrow |
| ACHMILLE | *Achillea millefolium* | Common Yarrow |
| ACOCALAM | *Acorus calamus* | Sweet-Flag |
| AGRCRIST | *Agropyron cristatum* ssp.<br>*pectinatum* | Crested Wheatgrass |
| AGRGIGAN | *Agrostis gigantea* | Redtop |
| AGRSCABR | *Agrostis scabra* | Ticklegrass |
| AGRSTRIA | *Agrimonia striata* | Roadside Agrimony |
| ALITRIVI | *Alisma triviale* | Northern Water Plantain |
| ALOEAEQUA | *Alopecurus aequalis* | Short-Awn Meadow-Foxtail |
| ALOPRATE | *Alopecurus pratensis* | Field Meadow-Foxtail |
| AMARETRO | *Amaranthus retroflexus* | Redroot Pigweed |
| AMEALNIF | *Amelanchier alnifolia* | Saskatoon Berry |
| ANAMINIM | *Anagallis minima* | Chaffweed |
| ANECANAD | *Anemone canadensis* | Canada Anemone |
| ANTPARVI | *Antennaria parvifolia* | Small-Leaf Pussytoes |
| ARNCHAMI | *Arnica chamissonis* | Chamisso Arnica |
| ARTBIENN | *Artemisia biennis* | Biennial Sagewort |
| ARTCAMPE | *Artemisia campestris* ssp. *caudata* | Common Sagewort |
| ARTLONGI | *Artemisia longifolia* | Longleaf Sagebrush |
| ARTLUDOV | *Artemisia ludoviciana* | Gray Sagewort |
| ATRPROST | *Atriplex prostrata* | Triangle Orache |
| AVEFATUA | *Avena fatua* | Wild Oats |
| BECSYZIG | *Beckmannia syzigachne* ssp.<br>*syzigachne* | American Sloughgrass |
| BIDCERNU | *Bidens cernua* | Nodding Beggarticks |
| BOLMARIT | *Bolboschoenus maritimus* | Cosmopolitan Bulrush |
| BRANAPUS | *Brassica napus* | Argentine Canola |
| BROINERM | *Bromus inermis* | Smooth Brome |
| CALCANAD | *Calamagrostis canadensis* var.<br>*canadensis* | Bluejoint |
| CALIPALU | *Callitriche palustris* | Vernal Water-Starwort |
| CALLPALU | *Calla palustris* | Water Arum |
| CALSTRIC | *Calamagrostis stricta* ssp. *inexpansa* | Slimstem Reedgrass |
| CALTPALU | *Caltha palustris* | Yellow Marsh Marigold |
| CAPBURSA | *Capsella bursa-pastoris* | Shepherd's Purse |
| CARAQUAT | *Carex aquatilis* | Water Sedge |
| CARATHER | *Carex atherodes* | Awed Sedge |
| CARATHRO | *Carex athrostachya* | Slender-Beak Sedge |
| CARBEBBI | *Carex bebbii* | Bebb's Sedge |
| CARBREVI | *Carex brevior* | Shortbeak Sedge |
| CARCARVI | *Carum carvi* | Wild Caraway |
| CARDIAND | *Carex diandra* | Two-Stamened Sedge |
| CAREX_SP | *Carex sp.* | Sedge |
| CARLACUS | *Carex lacustris* | Lakebank Sedge |
| CARPELLI | *Carex pellita* | Woolly Sedge |
| CARPRAEG | *Carex praegracilis* | Clustered Field Sedge |
| CARPRATI | *Carex praticola* | Meadow Sedge |
| CARRETRO | *Carex retrorsa* | Knotsheath Sedge |
| CARSARTW | *Carex sartwellii* | Sartwell's Sedge |
| CARSYCHN | *Carex sychnocephala* | Many-Headed Sedge |
| CARUTRIC | *Carex utriculata* | Northwest Territory Sedge |
| CERARVEN | *Cerastium arvense* | Field Chickweed |
| CHAANGUS | *Chamerion angustifolium ssp. angustifolium* | Fireweed |
| CHEALBUM | *Chenopodium album* | Common Lambsquarters |
| CHECAPIT | *Chenopodium capitatum* | Strawberry Blite |
| CHERUBRU | *Chenopodium rubrum* | Red Goosefoot |
| CICMACUL | *Cicuta maculata var. angustifolia* | Spotted Water Hemlock |
| CIRARVEN | *Cirsium arvense* | Canada Thistle |
| CIRVULGA | *Cirsium vulgare* | Bull Thistle |
| COLLINEA | *Collomia linearis* | Narrow-Leaf Mountain Trumpet |
| COMPALUS | *Comarum palustre* | Purple Marshlocks |
| CORSERIC | *Cornus sericea ssp. sericea* | Red Osier Dogwood |
| CRETECTO | *Crepis tectorum* | Narrow-Leaf Hawk's Beard |
| DESCESPI | *Deschampsia cespitosa ssp. cespitosa* | Tufted Hairgrass |
| DESSOPHI | *Descurainia sophia* | Flaxweed Tansymustard |
| ECHCRUSG | *Echinochloa crus-galli* | Barnyard Grass |
| ELACOMMU | *Elaeagnus commutata* | Silverberry |
| ELEACICU | *Eleocharis acicularis* | Needle Spikerush |
| ELEPALUS | *Eleocharis palustris* | Creeping Spikerush |
| ELYREPEN | *Elymus repens* | Quackgrass |
| ELYTRACH | *Elymus trachycaulus* | Slender Wheatgrass |
| EPICAMPE | *Epilobium campestre* | Smooth Spike-Primrose |
| EPICILIA | *Epilobium ciliatum ssp.* | Fringed Willow-Herb |
|  | *glandulosum* |  |
| EPILEPTO | *Epilobium leptophyllum* | Bog Willow-Herb |
| EIPALUS | *Epilobium palustre* | Marsh Willow-Herb |
| EQUARVEN | *Equisetum arvense* | Common Horsetail |
| EQUFLUVI | *Equisetum fluviatile* | River Horsetail |
| EQUHYMAL | *Equisetum hyemale ssp. affine* | Scouring Horsetail |
| EQUPALUS | *Equisetum palustre* | Marsh Horsetail |
| EQUPRATE | *Equisetum pratense* | Meadow Horsetail |
| ERIGRACI | *Eriophorum gracile* | Slender Cotton-Grass |
| ERILONCH | *Erigeron lonchophyllus* | Low-Meadow Fleabane |
| ERIPHILA | *Erigeron philadelphicus* | Philadelphia Fleabane |
| ERUGALLI | *Erucastrum gallicum* | Common Dog-Mustard |
| ERYCHEIR | *Erysimum cheiranthoides* | Wallflower Mustard |
| EURCONSP | *Eurybia conspicua* | Western Showy Aster |
| FAGESCUL | *Fagopyrum esculentum* | Common Buckwheat |
| FALCONVO | *Fallopia convolvulus* | Black Bindweed |
| FALSCAND | *Fallopia scandens* | Climbing False Buckwheat |
| FESSAXIM | *Festuca saximontana* | Rocky Mountain Fescue |
| FRAVESCA | *Fragaria vesca* | Woodland Strawberry |
| FRAVIRGI | *Fragaria virginiana ssp. glauca* | Wild Strawberry |
| GALTETRA | *Galeopsis tetrahit* | Brittle-Stem Hedge-Nettle |
| GALTRIFI | *Galium trifidum* | Small Bedstraw |
| GALTRIFL | *Galium triflorum* | Sweet Bedstraw |
| GEUALEPP | *Geum aleppicum* | Yellow Avens |
| GEUMACRO | *Geum macrophyllum var. princisum* | Large-Leaf Avens |
| GEURIVAL | *Geum rivale* | Purple Avens |
| GLYBOREA | *Glyceria borealis* | Northern Manna Grass |
| GLYGRAND | *Glyceria grandis* | American Manna Grass |
| GLYSTRIA | *Glyceria striata* | Fowl Manna Grass |
| GRANEGLE | *Gratiola neglecta* | Clammy Hedge-Hyssop |
| GRISQUAR | *Grindelia squarrosa* | Curlytop Gumweed |
| HIEUMBAL | *Hieracium umbellatum* | Canadian Hawkweed |
| HIPVULGA | *Hippuris vulgaris* | Common Mare's Tail |
| HORJUBAT | *Hordeum jubatum* | Foxtail Barley |
| HORVULGA | *Hordeum vulgare* | Common Barley |
| JUNBALTI | *Juncus balticus ssp. ater* | Baltic Rush |
| JUNLONGI | *Juncus longistylus* | Long-Style Rush |
| JUNNODOS | *Juncus nodosus* | Jointed Rush |
| JUNVASEY | *Juncus vaseyi* | Vasey's Rush |
| KRALANAT | *Krascheninnikovia lanata* | Winterfat |
| LACSERRI | *Lactuca serriola* | Prickly Lettuce |
| LATOCHRO | *Lathyrus othroleucus* | Cream Peavine |
| LEMMINOR | *Lemna minor* | Common Duckweed |
| LEMTRISU | *Lemna trisulca* | Ivy-Leaf Duckweed |
| LINUSITA | *Linum usitatissimum* | Common Flax |
| LYCASPER | *Lycopus asper* | Rough Water Hore-Hound |
| LYSCILIA | *Lysimachia ciliata* | Fringed Yellow Loosestrife |
| LYSMARIT | *Lysimachia maritima* | Sea Milkwort |
| LYSTHYRS | *Lysimachia thyrsiflora* | Tufted Yellow Loosestrife |
| MAISTELL | *Maianthemum stellatum* | False Solomon's Seal |
| MALNEGLE | *Malva neglecta* | Roundleaf Mallow |
| MEDSATIV | *Medicago sativa* | Alfalfa |
| MELALBUS | *Melilotus albus* | White Sweet-Clover |
| MELIL_SP | *Melilotus sp.* | Sweet-Clover |
| MENARVEN | *Mentha arvensis* | Wild Mint |
| MONNUTTA | *Monolepis nuttalliana* | Nuttall's Poverty-Weed |
| MUHRICHA | *Muhlenbergia richardsonis* | Mat Muhly |
| MULOBLON | *Mulgedium oblongifolium* | Blue Lettuce |
| PENPROCE | *Penstemon procerus* | Pincushion Beardtongue |
| PERAMPHI | *Persicaria amphibia* | Water Knotweed |
| PERLAPAT | *Persicaria lapathifolia* | Curlytop Knotweed |
| PETFRIGI | *Petasites frigidus var. sagittatus* | Arctic Sweet Colt's-Foot |
| PHAARUND | *Phalaris arundinacea* | Reed Canary Grass |
| PHLPRATE | *Phleum pratense* | Common Timothy |
| PLAHYPER | *Platanthera hyperborea* | Northern Bog Orchid |
| PLAMAJOR | *Plantago major* | Broadleaf Plantain |
| PLASCOUL | *Plagiobothrys scouleri* | Scouler's Popcornflower |
| POAPALUS | *Poa palustris* | Fowl Bluegrass |
| POAPRATE | *Poa pratensis* | Kentucky Bluegrass |
| POLAVICU | *Polygonum aviculare ssp. depressum* | Prostrate Knotweed |
| POLRAMOS | *Polygonum ramosissimum* | Bushy Knotweed |
| POPBALSA | *Populus balsamifera* | Balsam Poplar |
| POPTREMU | *Populus tremuloides* | Trembling Aspen |
| POTANSER | *Potentilla anserina* | Silverweed Cinquefoil |
| POTGRAMI | *Potamogeton gramineus* | Variableleaf Pondweed |
| POTNORVE | *Potentilla norvegica* | Norwegian Cinquefoil |
| POTRICHA | *Potamogeton richardsonii* | Richardson's Pondweed |
| POTRIVAL | *Potentilla rivalis* | Brook Cinquefoil |
| PYRASARI | *Pyrola asarifolia* | Pink Wintergreen |
| RANAQUAT | *Ranunculus aquatilis* var. *diffusus* | Water Buttercup |
| RANCYMBA | *Ranunculus cymbalaria* | Alkali Buttercup |
| RANGMELI | *Ranunculus gmelinii* | Gmelin's Buttercup |
| RANMACOU | *Ranunculus macounii* | Macoun's Buttercup |
| RANSCELE | *Ranunculus sceleratus* var. *multifidus* | Celeryleaf Buttercup |
| RANUN_SP | *Ranunculus* sp. | Buttercup |
| RIBLACUS | *Ribes lacustre* | Prickly Currant |
| RIBOXYAC | *Ribes oxycanthoides* | Canadian Gooseberry |
| RORPALUS | *Rorippa palustris* | Marsh Yellowcress |
| ROSACICU | *Rosa acicularis* ssp. *sayi* | Prickly Rose |
| RUBPUBES | *Rubus pubescens* | Dwarf Red Raspberry |
| RUBSACHA | *Rubus sachalinensis* var. *sachalinensis* | Common Red Raspberry |
| RUMBRITA | *Rumex britannica* | Greater Water Dock |
| RUMCRISP | *Rumex crispus* | Curly Dock |
| RUMEX_SP | *Rumex* sp. | Dock |
| RUMFUEGI | *Rumex fueginus* | Golden Dock |
| RUMOCCID | *Rumex occidentalis* | Western Dock |
| RUMSALIC | *Rumex salicifolius* | Willow Dock |
| SAGCUNEA | *Sagittaria cuneata* | Arum-Leaf Arrowhead |
| SALBEBBI | *Salix bebbiana* | Bebb's Willow |
| SALDISCO | *Salix discolor* | Pussy Willow |
| SALEXIGU | *Salix exigua* | Sandbar Willow |
| SALIX_SP | *Salix* sp. | Willow |
| SALLASIA | *Salix lasiandra* var. *lasiandra* | Pacific Willow |
| SALLUCID | *Salix lucida* | Shining Willow |
| SALMACCA | *Salix maccalliana* | McCalla's Willow |
| SALPETIO | *Salix petiolaris* | Meadow Willow |
| SALPLANI | *Salix planifolia* | Plain-Leaf Willow |
| SALPSEUD | *Salix pseudomonticola* | False Mountain Willow |
| SALPYRIF | *Salix pyrifolia* | Balsam Willow |
| SALRUBRA | *Salicornia rubra* | Red Samphire |
| SALSERIS | *Salix serissima* | Autumn Willow |
| SCHACUTU | *Schoenoplectus acutus* var. *acutus* | Hard-Stem Bulrush |
| SCHOE_SP | *Schoenoplectus* sp. | Bulrush |
| SCHPUNGE | *Schoenoplectus pungens* var. | Common Three-Square Bulrush |
|  | *pungens* |  |
| SCHTABER | *Schoenoplectus tabernaemontani* | Soft-Stem Bulrush |
| SCIMICRO | *Scirpus microcarpus* | Panicked Bulrush |
| SCOFESTU | *Scolochloa festucacea* | Common Rivergrass |
| SCUGALER | *Scutellaria galericulata* | Marsh Skullcap |
| Sd_Forb | - | Unidentifiable seedling/forb |
| SENVULGA | *Senecio vulgaris* | Common Groundsel |
| SISMONTA | *Sisyrinchium montanum* | Mountain Blue-Eyed Grass |
| SIUSUAVE | *Sium suave* | Common Water Parsnip |
| SOLALTIS | *Solidago altissima ssp. gilvocanescens* | Canada Goldenrod |
| SONARVEN | *Sonchus arvensis* | Perennial Sow-Thistle |
| SONASPER | *Sonchus asper* | Prickly Sow-Thistle |
| SONOLERA | *Sonchus oleraceus* | Annual Sow-Thistle |
| SPAANGUS | *Sparganium angustifolium* | Narrow-Leaf Bur-Reed |
| SPAEURYC | *Sparganium eurycarpum* | Giant Bur-Reed |
| SPESALIN | *Spergularia salina* | Salt Sandspurry |
| SPHINTER | *Sphenopholis intermedia* | Slender Wedgegrass |
| SPOCRYPT | *Sporobolus cryptandrus* | Sand Dropseed |
| STAPILOS | *Stachys pilosa var. pilosa* | Hairy Hedgenettle |
| STELONGI | *Stellaria longifolia* | Long-Leag Starwort |
| STEMEDIA | *Stellaria media* | Common Chickweed |
| SUACALCE | *Suaeda calceoliformis* | Paiuteweed |
| SYMBOREA | *Symphyotrichum boreale* | Northern Bog Aster |
| SYMERICO | *Symphyotrichum ericoides var. pansum* | White Heath Aster |
| SYMLANCE | *Symphyotrichum lanceolatum var. hesperium* | White Panicle Aster |
| SYMOCCID | *Symphoricarpos occidentalis* | Western Snowberry |
| SYMPUNIC | *Symphyotrichum puniceum var. puniceum* | Purplestem Aster |
| TANVULGA | *Tanacetum vulgare* | Common Tansy |
| TAROFFIC | *Taraxacum officinale* | Common Dandelion |
| TEPPALUS | *Tephrosia palustris* | Marsh Fleabane |
| THLARVEN | *Thlaspi arvense* | Field Pennycress |
| TRADUBIU | *Tragopogon dubius* | Goat's Beard |
| TRIHYBRI | *Trifolium hybridum* | Alsike Clover |
| TRIMARIT | *Triglochin maritima* | Seaside Arrow-Grass |
| TYPLATIF | *Typha latifolia* | Common Cattail |
| UNKASTER | *Asteraceae* | Unidentifiable Asteraceae |

| **Code** | **Scientific Name** | **Common Name** |
| --- | --- | --- |
| UNKBRASS | *Brassicaceae* | Unidentifiable Brassicaceae |
| UNKCARYO | *Caryophyllaceae* | Unidentifiable Caryophyllaceae |
| UNKPOACE | *Poaceae* | Unidentifiable Poaceae |
| URTDIOCA | *Urtica dioica ssp. gracilis* | Stinging Nettle |
| UTRVULGA | *Utricularia vulgaris ssp. macrorhiza* | Common Bladderwort |
| VERPEREG | *Veronica peregrina* | Purslane Speedwell |
| VERSCUTE | *Veronica scutellata* | Marsh Speedwell |
| VICAMERI | *Vicia americana* | American Vetch |
| VIOADUNC | *Viola adunca* | Early Blue Violet |
| VIOCANAD | *Viola canadensis* | Canada White Violet |
| VIOSOROR | *Viola sororia var. affinis* | Common Blue Violet |

**Appendix S2A.** List of bird functional traits; traits values are reported in Anderson (2017)^1^.

| **Group Class** | **Group Code** | **Meaning** |
| --- | --- | --- |
| Diet Classification | Carniv | Carnivore |
| Diet Classification | Herbiv | Herbivore |
| Diet Classification | Omniv | Omnivore |
| Migration Habitat | Res | Resident |
| Migration Habitat | TrpclMgr | Tropical Migrant |
| Primary Diet | AqInsect | Aquatic Insects |
| Primary Diet | AqPlnts | Aquatic Plants |
| Primary Diet | Carrion | Carrion |
| Primary Diet | Fish | Fish |
| Primary Diet | Fruit | Fruit |
| Primary Diet | Grains | Grains |
| Primary Diet | Insects | Insects |
| Primary Diet | Nuts | Nuts |
| Primary Diet | Plants | Plants |
| Primary Diet | Seeds | Seeds |
| Primary Diet | SmAnml | Small Animals |
| Primary Feeding Habit | ArlFrgr | Aerial Forager |
| Primary Feeding Habit | ArlPrst | Aerial Pursuit |
| Primary Feeding Habit | BrkGln | Bark Gleaner |
| Primary Feeding Habit | Dbblr | Dabbler |
| Primary Feeding Habit | FlgGln | Foliage Gleaner |
| Primary Feeding Habit | GrndFrg | Ground Forager |
| Primary Feeding Habit | GrndGln | Ground Gleaner |
| Primary Feeding Habit | HPatrol | Hawk and Patrol |
| Primary Feeding Habit | HvrGln | Hovers and Gleaners |
| Primary Feeding Habit | Insect | Insectivore |
| Primary Feeding Habit | Prbs | Probbing |
| Primary Feeding Habit | SrfcDvr | Surface Diver |
| Primary Feeding Habit | Stlknng | Stalking |
| Primary Habitat | Field | Field |
| Primary Habitat | Forest | Forest |
| Primary Habitat | Grsslnd | Grassland |
| Primary Habitat | LkPd | Lake or Pond |
| Primary Habitat | Marsh | Marsh |
| Primary Habitat | OpnWood | Open Woodland |
| Primary Habitat | RvrStrm | River or Stream |
| Primary Habitat | Scrub | Scrub |
| Primary Habitat | Shrln | Shoreline |
| Primary Nesting Location | Bank | Bank |

<sup>1</sup> Anderson, D. 2017. Monitoring wetland integrity and restoration success with avifauna in the Prairie Pothole Region of Alberta, Canada. University of Waterloo

| Group Class | Group Code | Meaning |
| --- | --- | --- |
| Primary Nesting Location | Cavity | Cavity |
| Primary Nesting Location | Fltng | Floating |
| Primary Nesting Location | Grnd | Ground |
| Primary Nesting Location | Reeds | Reeds |
| Primary Nesting Location | Shrb | Shrub |
| Primary Nesting Location | Strctr | Structure |
| Primary Nesting Location | Tree | Tree |
| Wetland Status | Facul | Facultative |
| Wetland Status | FaculDry | Facultative Dry |
| Wetland Status | FaculWet | Facultative Wet |

**Appendix S2B.** List of aquatic macroinvertebrates functional traits; traits values are reported in Gleason (2017)^2^.

| Group Class | Group Code | Meaning |
| --- | --- | --- |
| Feeding Groups | ENGULF | Engulfing Predators |
| Feeding Groups | FCOLL | Filtering Collectors |
| Feeding Groups | GCOLL | Gathering Collectors |
| Feeding Groups | SCRAPE | Scrapers |
| Feeding Groups | SHRED | Shredders |
| Feeding Groups | PARA | Ecotoparasites |
| Feeding Groups | PIERCE | Piercing Predators |
| Behavioural Guilds | BUR | Burrowers |
| Behavioural Guilds | CLIMB | Climber |
| Behavioural Guilds | CLING | Clinger |
| Behavioural Guilds | DIVER | Diver |
| Behavioural Guilds | SKATE | Skater |
| Behavioural Guilds | SPRAWLER | Sprawler |
| Behavioural Guilds | SWIM | Swimmer |
| Desiccation Strategy Groups | TOLE | Tolerators |
| Desiccation Strategy Groups | WETL | Wet Layers |
| Desiccation Strategy Groups | DRYL | Dry Layers |
| Desiccation Strategy Groups | DISP | Dispersers |

<sup>2</sup> Gleason, J. E. 2017. Aquatic macroinvertebrate communities and diversity patterns in the Northern Prairie Pothole Region. University of Waterloo.

**Appendix S2C.** List of plant functional traits; traits values are reported in Bolding (2018)^3^.

| **Group Class** | **Group Code** | **Meaning** |
| --- | --- | --- |
| Wetland Plant Indicator Status | Forb | Forb |
| Wetland Plant Indicator Status | Graminoid | Graminoid |
| Wetland Plant Indicator Status | Vine | Vine |
| Wetland Plant Indicator Status | Hardwood | Hardwood |
| Wetland Plant Indicator Status | Tall_Shrub | Tall Shrub |
| Wetland Plant Indicator Status | Low_Shrub | Low Shrub |
| Wetland Plant Indicator Status | BroadLeaf_Emergent | BroadLeaf Emergent |
| Wetland Plant Indicator Status | Floating_Plant | Floating Plant |
| Wetland Plant Indicator Status | FreeFloating_Plant | FreeFloating Plant |
| Wetland Plant Indicator Status | NarrowLeaf_Emergent | NarrowLeaf Emergent |
| Wetland Plant Indicator Status | Robust_Emergent | Robust Emergent |
| Wetland Plant Indicator Status | Submersed_Plant | Submersed Plant |
| Vegetative Reproduction | Vegetative_Reproduction | Vegetative Reproduction |
| Nitrogen Fixing | Nitrogen_Fixer | Nitrogen Fixer |
| Litter Decomposal | Recalcitrant_Litter | Recalcitrant Litter |
| Native Status | Native_Graminoid | Native Graminoid |
| Native Status | Exotic_Graminoid | Exotic Graminoid |
| Native Status | Native_Perennials | Native Perennials |
| Native Status | Exotic_Perennials | Exotic Perennials |
| Native Status | Native_Annuals_Biennials | Native Annuals Biennials |
| Native Status | Exotic_Annuals_Biennials | Exotic Annuals and Biennials |
| Dispersal Mechanism | Anemochory | Anemochory |
| Dispersal Mechanism | Hydrochory | Hydrochory |
| Dispersal Mechanism | Zoochory | Zoochory |
| Dispersal Mechanism | Multiple_Dispersal | Multiple Dispersal |

## Footnotes

1 Anderson, D. 2017. Monitoring wetland integrity and restoration success with avifauna in the Prairie Pothole Region of Alberta, Canada. University of Waterloo

2 Gleason, J. E. 2017. Aquatic macroinvertebrate communities and diversity patterns in the Northern Prairie Pothole Region. University of Waterloo

3 Bolding, M. 2018. Vegetation based assessment of wetland condition in the Prairie Pothole Region. University of Waterloo.

## Notes

### Competing Interest Statement

The authors have declared no competing interest.

## References

Ackerly, D. D., and W. K. Cornwell. 2007. A trait-based approach to community assembly: partitioning of species trait values into within- and among-community components. Ecology Letters 10:135–145.

Anderson, D. L., and R. C. Rooney. 2019. Differences exist in bird communities using restored and natural wetlands in the Parkland Region, Alberta, Canada. Restoration Ecology 27:1495–1507.

Aronson, M. F. J., C. H. Nilon, C. A. Lepczyk, T. S. Parker, P. S. Warren, S. S. Cilliers, M. A. Goddard, A. K. Hahs, C. Herzog, M. Katti, F. A. La Sorte, N. S. G. Williams, and W. Zipperer. 2016. Hierarchical filters determine community assembly of urban species pools. Ecology 97:2952–2963.

Austin, J. E., and D. A. Buhl. 2011. Nest survival of American Coots relative to grazing, burning, and water depths. Avian Conservation and Ecology 6: art1. http://dx.doi.org/10.5751/ACE-00472-060201

Ayers, C. R., K. C. Hanson-Dorr, S. O’Dell, C. D. Lovell, M. L. Jones, J. R. Suckow, and B. S. Dorr. 2015. Impacts of colonial waterbirds on vegetation and potential restoration of island habitats. Restoration Ecology 23:252–260.

Azeria, E. T., D. Fortin, J. Lemaître, P. Janssen, C. Hébert, M. Darveau, and S. G. Cumming. 2009. Fine-scale structure and cross-taxon congruence of bird and beetle assemblages in an old-growth boreal forest mosaic. Global Ecology and Biogeography 18:333–345.

Bolding, M. T., A. J. Kraft, D. T. Robinson, and R. C. Rooney. 2020. Improvements in multi-metric index development using a whole-index approach. Ecological Indicators 113:106191.

Bowman, W. D., T. A. Theodose, J. C. Schardt, and R. T. Conant. 1993. Constraints of nutrient availability on primary production in two alpine tundra communities. Ecology 74:2085–2097.

Cabra-García, J., C. Bermúdez-Rivas, A. M. Osorio, and P. Chacón. 2012. Cross-taxon congruence of α and β diversity among five leaf litter arthropod groups in Colombia. Biodiversity and Conservation 21:1493–1508.

Campeau, S., H. R. Murkin, and R. D. Titman. 1994. Relative importance of algae and emergent plant litter to freshwater marsh invertebrates. Canadian Journal of Fisheries and Aquatic Sciences 51:681–692.

Casanova, M. T., and M. A. Brock. 2000. How do depth, duration and frequency of flooding influence the establishment of wetland plant communities? Plant Ecology 147:237–250.

Chase, J. M., and J. A. Myers. 2011. Disentangling the importance of ecological niches from stochastic processes across scales. Philosophical Transactions of the Royal Society B: Biological Sciences 366:2351–2363.

Clifford, H. F. 1991. Aquatic Invertebrates of Alberta. University of Alberta Press, Edmonton, Alberta.

Comte, L., J. Cucherousset, S. Boulêtreau, and J. D. Olden. 2016. Resource partitioning and functional diversity of worldwide freshwater fish communities. Ecosphere 7: e01356.

Connor, E. F., and D. Simberloff. 1979. The assembly of species communities: chance or competition. Ecology 60:1132.

Daniel, J., J. E. Gleason, K. Cottenie, and R. C. Rooney. 2019. Stochastic and deterministic processes drive wetland community assembly across a gradient of environmental filtering. Oikos 128:1158–1169.

Davis, C. A., and J. R. Bidwell. 2008. Response of aquatic invertebrates to vegetation management and agriculture. Wetlands 28:793–805.

Devercelli, M., P. Scarabotti, G. Mayora, B. Schneider, and F. Giri. 2016. Unravelling the role of determinism and stochasticity in structuring the phytoplanktonic metacommunity of the Paraná River floodplain. Hydrobiologia 764:139–156.

Diamond, A. W. 1975. Assembly of species communities. Page in M. L. Cody and J. M. Diamond, editors. Ecology and Evolution of Communities. Harvard University Press, Cambridge, Massachusetts.

Dormann, C. F., M. Bobrowski, D. M. Dehling, D. J. Harris, F. Hartig, H. Lischke, M. D. Moretti, J. Pagel, S. Pinkert, M. Schleuning, S. I. Schmidt, C. S. Sheppard, M. J. Steinbauer, D. Zeuss, and C. Kraan. 2018. Biotic interactions in species distribution modelling: 10 questions to guide interpretation and avoid false conclusions. Global Ecology and Biogeography 27:1004–1016.

Downing, D. J., and W. W. Pettapiece. 2006. Natural Regions and Subregions of Alberta. Government of Alberta, Edmonton, Alberta.

Duan, M., Y. Liu, Z. Yu, J. Baudry, L. Li, C. Wang, and J. C. Axmacher. 2016. Disentangling effects of abiotic factors and biotic interactions on cross-taxon congruence in species turnover patterns of plants, moths and beetles. Scientific Reports 6:23511.

Euliss, N. H., J. W. Labaugh, L. H. Fredrickson, D. M. Mushet, M. K. Laubhan, G. A. Swanson, T. C. Winter, D. O. Rosenberry, and R. D. Nelson. 2004. The wetland continuum: a conceptual framework for interpreting biological studies. Wetlands 24:448–458.

Environment Canada, 2014. CABIN laboratory methods: processing, taxonomy, and quality control of benthic macroinvertebrate samples. http://publications.gc.ca/collections/collection_2015/ec/En84-86-2014-eng.pdf

Finke, D. L., and W. E. Snyder. 2008. Niche partitioning increases resource exploitation by diverse communities. Science 321:1488–1490.

Fourqurean, J. W., J. C. Zieman, and G. V. N. Powell. 1992. Phosphorus limitation of primary production in Florida Bay: Evidence from C: N: P ratios of the dominant seagrass *Thalassia testudinum*. Limnology and Oceanography 37:162–171.

Fox, A. D., L. Cao, Y. Zhang, M. Barter, M. J. Zhao, F. J. Meng, and S. L. Wang. 2011. Declines in the tuber-feeding waterbird guild at Shengjin Lake National Nature Reserve, China – a barometer of submerged macrophyte collapse. Aquatic Conservation: Marine and Freshwater Ecosystems 21:82–91.

Galatowitsch, S. M. 2006. Restoring prairie pothole wetlands: does the species pool concept offer decision-making guidance for re-vegetation? Applied Vegetation Science 9:261–270.

Gallardo, L. I., R. P. Carnevali, E. A. Porcel, and A. S. G. Poi. 2011. Does the effect of aquatic plant types on invertebrate assemblages change across seasons in a subtropical wetland? Limnetica 29:87–98.

García, D. 2016. Birds in ecological networks: insights from bird-plant mutualistic interactions. Ardeola 63:151–180.

Gerhold, P., J. F. Cahill, M. Winter, I. V. Bartish, and A. Prinzing. 2015. Phylogenetic patterns are not proxies of community assembly mechanisms (they are far better). Functional Ecology 29:600–614.

Gleason, J. E., J. Y. Bortolotti, and R. C. Rooney. 2018. Wetland microhabitats support distinct communities of aquatic macroinvertebrates. Journal of Freshwater Ecology 33:73–82.

Gleason, J. E., and R. C. Rooney. 2017. Aquatic macroinvertebrates are poor indicators of agricultural activity in northern prairie pothole wetlands. Ecological Indicators 81:333–339.

Gleason, J. E., and R. C. Rooney. 2018. Pond permanence is a key determinant of aquatic macroinvertebrate community structure in wetlands. Freshwater Biology 63:264–277.

Gotelli, N. J. 2000. Null model analysis of species co-occurrence patterns. Ecology 81:2606.

Gotelli, N. J., and D. J. McCabe. 2002. Species co-occurrence: a meta-analysis of J. M. Diamond’s assembly rules model. Ecology 83:2091–2096.

Gotelli, N. J., and K. Rohde. 2002. Co-occurrence of ectoparasites of marine fishes: a null model analysis. Ecology Letters 5:86–94.

Government of Alberta. 2014. Alberta Merged Wetland Inventory. Alberta Environment and Parks, Government of Alberta, Edmonton, Alberta.

Grace, J. B., T. M. Anderson, H. Olff, and S. M. Scheiner. 2010. On the specification of structural equation models for ecological systems. Ecological Monographs 80:67–87.

Groendahl, S., and P. Fink. 2017. Consumer species richness and nutrients interact in determining producer diversity. Scientific Reports 7:44869.

Guarino, R., B. Ferrario, and L. Mossa. 2005. A stochastic model of seed dispersal pattern to assess seed predation by ants in annual dry grasslands. Plant Ecology 178:225–235.

Guignard, M. S., A. R. Leitch, C. Acquisti, C. Eizaguirre, J. J. Elser, D. O. Hessen, P. D. Jeyasingh, M. Neiman, A. E. Richardson, P. S. Soltis, D. E. Soltis, C. J. Stevens, M. Trimmer, L. J. Weider, G. Woodward, and I. J. Leitch. 2017. Impacts of nitrogen and phosphorus: from genomes to natural ecosystems and agriculture. Frontiers in Ecology and Evolution 5:70.

Gurney, K. E. B., R. G. Clark, S. M. Slattery, and L. C. M. Ross. 2017. Connecting the trophic dots: responses of an aquatic bird species to variable abundance of macroinvertebrates in northern boreal wetlands. Hydrobiologia 785:1–17.

Hall, D. L., M. R. Willig, D. L. Moorhead, R. W. Sites, E. B. Fish, and T. R. Mollhagen. 2004. Aquatic macroinvertebrate diversity of playa wetlands: The role of landscape and island biogeographic characteristics. Wetlands 24:77–91.

Hayashi, M., G. van der Kamp, and D. O. Rosenberry. 2016. Hydrology of prairie wetlands: understanding the integrated surface-water and groundwater processes. Wetlands 36:237–254.

Horváth, Z., M. Ferenczi, A. Móra, C. F. Vad, A. Ambrus, L. Forró, G. Szövényi, and S. Andrikovics. 2012. Invertebrate food sources for waterbirds provided by the reconstructed wetland of Nyirkai-Hany, northwestern Hungary. Hydrobiologia 697:59–72.

Hurtt, G.C. and S.W. Pacala. 1995. The consequences of recruitment limitation: reconciling chance, history and competitive differences between plants. Journal of Theoretical Biology 176:1–12.

Jackson, A. C., M. G. Chapman, and A. J. Underwood. 2008. Ecological interactions in the provision of habitat by urban development: whelks and engineering by oysters on artificial seawalls. Austral Ecology 33:307–316.

Jorgensen, T. D., S. Pornprasertmanit, A. M. Schoemann, and Y. Rosseel. 2018. semTools: useful tools for structural equation modeling. R package version 0.5-1.

Kantrud, H. a, and R. E. Stewart. 1984. Ecological distribution and crude density of breeding birds on prairie wetlands. The Journal of Wildlife Management 48:426.

Klaassen, M., and B. A. Nolet. 2007. The role of herbivorous water birds in aquatic systems through interactions with aquatic macrophytes, with special reference to the Bewick’s Swan – Fennel Pondweed system. Hydrobiologia 584:205–213.

Kraft, A. J., D. T. Robinson, I. S. Evans, and R. C. Rooney. 2019. Concordance in wetland physicochemical conditions, vegetation, and surrounding land cover is robust to data extraction approach. PLOS ONE 14: e0216343.

Kraft, N. J. B., P. B. Adler, O. Godoy, E. C. James, S. Fuller, and J. M. Levine. 2015. Community assembly, coexistence and the environmental filtering metaphor. Functional Ecology 29:592–599.

LaBaugh, J. W., D. O. Rosenberry, D. M. Mushet, B. P. Neff, R. D. Nelson, and N. H. Euliss. 2018. Long-term changes in pond permanence, size, and salinity in Prairie Pothole Region wetlands: The role of groundwater-pond interaction. Journal of Hydrology: Regional Studies 17:1–23.

Laliberte, E., P. Legendre, and B. Shipley. 2014. FD: measuring functional diversity from multiple traits, and other tools for functional ecology. R package version 1.3.

Lee Foote, A., and C. L. Rice Hornung. 2005. Odonates as biological indicators of grazing effects on Canadian prairie wetlands. Ecological Entomology 30:273–283.

van Leeuwen, C. H. A., Á. Lovas-Kiss, M. Ovegård, and A. J. Green. 2017. Great cormorants reveal overlooked secondary dispersal of plants and invertebrates by piscivorous waterbirds. Biology Letters 13:20170406.

Leibowitz, S. G., and K. C. Vining. 2003. Temporal connectivity in a prairie pothole complex. Wetlands 23:13–25.

Lokemoen, J. T., and R. O. Woodward. 1992. Nesting waterfowl and water birds on natural islands in the Dakotas and Montana. Wildlife Society Bulletin 20:163–171.

Longcore, J. R., D. G. McAuley, G. W. Pendelton, C. R. Bennatti, T. M. Mingo, and K. L. Stromborg. 2006. Macroinvertebrate abundance, water chemistry, and wetland characteristics affect use of wetlands by avian species in Maine. Hydrobiologia 567:143–167.

Maestre, F. T., C. Escolar, I. Martínez, and A. Escudero. 2008. Are soil lichen communities structured by biotic interactions? A null model analysis. Journal of Vegetation Science 19:261–266.

Magnuson, J. J., L. B. Crowder, and P. A. Medvick. 1979. Temperature as an ecological resource. American Zoologist 19:331–343.

Maurer, K. M., T. W. Stewart, and F. O. Lorenz. 2014. Direct and indirect effects of fish on invertebrates and tiger salamanders in prairie pothole wetlands. Wetlands 34:735–745.

Maynard, D. S., K. R. Covey, T. W. Crowther, N. W. Sokol, E. W. Morrison, S. D. Frey, L. T. A. van Diepen, and M. A. Bradford. 2018. Species associations overwhelm abiotic conditions to dictate the structure and function of wood-decay fungal communities. Ecology 99:801–811.

Merrit, R. W., K. W. Cummins, and M. B. Berg. 2008. An introduction to the aquatic insects of North America. Fourth edition. Kendall Hunt Publishing Company, Dubuque, Iowa.

Meyer, M. D., C. A. Davis, and J. R. Bidwell. 2013. Assessment of two methods for sampling invertebrates in shallow vegetated wetlands. Wetlands 33:1063–1073.

Meyer, M. D., C. A. Davis, and D. Dvorett. 2015. Response of wetland invertebrate communities to local and landscape factors in North Central Oklahoma. Wetlands 35:533–546.

Meyers, N. 2018. Use of water isotope tracers to characterize the hydrology of prairie wetlands in Alberta. Thesis. University of Waterloo, Waterloo, Ontario, Canada.

Morissette, J. L., K. J. Kardynal, E. M. Bayne, and K. A. Hobson. 2013. Comparing bird community composition among boreal wetlands: is wetland classification a missing piece of the habitat puzzle? Wetlands 33:653–665.

Mushet, D. M., O. P. McKenna, J. W. LaBaugh, N. H. Euliss, and D. O. Rosenberry. 2018. Accommodating state shifts within the conceptual framework of the wetland continuum. Wetlands 38:647–651.

Niemuth, N. D., M. E. Estey, R. E. Reynolds, C. R. Loesch, and W. A. Meeks. 2006. Use of wetlands by spring-migrant shorebirds in agricultural landscapes of North Dakota’s Drift Prairie. Wetlands 26:30–39.

O’Neal, B. J., E. J. Heske, and J. D. Stafford. 2008. Waterbird response to wetlands restored through the conservation reserve enhancement program. Journal of Wildlife Management 72:654–664.

Palacio, F. X., and J. M. Girini. 2018. Biotic interactions in species distribution models enhance model performance and shed light on natural history of rare birds: a case study using the straight-billed reedhaunter *Limnoctites rectirostris*. Journal of Avian Biology 49: e01743.

Poff, N. L. 1997. Landscape filters and species traits: towards mechanistic understanding and prediction in stream ecology. Journal of the North American Benthological Society 16:391–409.

Qian, H., and W. D. Kissling. 2010. Spatial scale and cross-taxon congruence of terrestrial vertebrate and vascular plant species richness in China. Ecology 91:1172–1183.

R Core Team. 2019. R: A language and environment for statistical computing. R Foundation for Statistical Computing, Vienna, Austria.

Rao, C. R. 1982. Diversity and dissimilarity coefficients: A unified approach. Theoretical Population Biology 21:24–43.

Reid, A. H., and W. G. Sprules. 2018. A comprehensive evaluation of *Daphnia pulex* foraging energetics and the influence of spatially heterogeneous food. Inland Waters 8:50–59.

Rosseel, Y. 2012. lavaan: an R package for structural equation modeling. Journal of Statistical Software 48:2.

Roy, M.-C., E. T. Azeria, D. Locky, and J. J. Gibson. 2019. Plant functional traits as indicator of the ecological condition of wetlands in the Grassland and Parkland of Alberta, Canada. Ecological Indicators 98:483–491.

Ruhí, A., E. Chappuis, D. Escoriza, M. Jover, J. Sala, D. Boix, S. Gascón, and E. Gacia. 2014. Environmental filtering determines community patterns in temporary wetlands: a multi-taxon approach. Hydrobiologia 723:25–39.

Schleuter, D., M. Daufresne, F. Massol, and C. Argillier. 2010. A user’s guide to functional diversity indices. Ecological Monographs 80:469–484.

Schoo, K. L., N. Aberle, A. M. Malzahn, and M. Boersma. 2012. Food quality affects secondary consumers even at low quantities: an experimental test with larval European Lobster. PLoS ONE 7: e33550.

Soons, M. B., A.-L. Brochet, E. Kleyheeg, and A. J. Green. 2016. Seed dispersal by dabbling ducks: an overlooked dispersal pathway for a broad spectrum of plant species. Journal of Ecology 104:443–455.

Spasojevic, M. J., and K. N. Suding. 2012. Inferring community assembly mechanisms from functional diversity patterns: the importance of multiple assembly processes. Journal of Ecology 100:652–661.

Spivak, A. C., E. A. Canuel, J. E. Duffy, and J. P. Richardson. 2009. Nutrient enrichment and food web composition affect ecosystem metabolism in an experimental seagrass habitat. PLoS ONE 4: e7473.

Stewart, R. E., and H. A. Kantrud. 1971. Classification of natural ponds and lakes in the glaciated prairie region. Page Bureau of Sport Fisheries and Wildlife Resource Publication 92. Washington, DC.

Thompson, R. C., B. J. Wilson, M. L. Tobin, A. S. Hill, and S. J. Hawkins. 1996. Biologically generated habitat provision and diversity of rocky shore organisms at a hierarchy of spatial scales. Journal of Experimental Marine Biology and Ecology 202:73–84.

Tiunov, A. V., and S. Scheu. 2005. Facilitative interactions rather than resource partitioning drive diversity-functioning relationships in laboratory fungal communities. Ecology Letters 8:618–625.

Tsai, J.-S., L. S. Venne, S. T. McMurry, and L. M. Smith. 2012. Local and landscape influences on plant communities in playa wetlands. Journal of Applied Ecology 49:174–181.

Urban, M. C., G. Bocedi, A. P. Hendry, J.-B. Mihoub, G. Peer, A. Singer, J. R. Bridle, L. G. Crozier, L. De Meester, W. Godsoe, A. Gonzalez, J. J. Hellmann, R. D. Holt, A. Huth, K. Johst, C. B. Krug, P. W. Leadley, S. C. F. Palmer, J. H. Pantel, A. Schmitz, P. A. Zollner, and J. M. J. Travis. 2016. Improving the forecast for biodiversity under climate change. Science 353: aad8466–aad8466.

van der Valk, A. G. 1981. Succession in wetlands: a Gleasonian approach. Ecology 62:688–696.

Vanausdall, R. A., and S. J. Dinsmore. 2019. Habitat associations of migratory waterbirds using restored shallow lakes in Iowa. Waterbirds 42:135.

Wagenmakers, E.-J., and S. Farrell. 2004. AIC model selection using Akaike weights. Psychonomic Bulletin & Review 11:192–196.

Wardle, D. A. 2006. The influence of biotic interactions on soil biodiversity. Ecology Letters 9:870–886.

Williams, N. S. G., M. W. Schwartz, P. A. Vesk, M. A. McCarthy, A. K. Hahs, S. E. Clemants, R. T. Corlett, R. P. Duncan, B. A. Norton, K. Thompson, and M. J. McDonnell. 2009. A conceptual framework for predicting the effects of urban environments on floras. Journal of Ecology 97:4–9.

Wright, H. E. J. 1972. Quaternary history of Minnesota. Pages 515–546 in P. K. Sims and G. Morey, editors. Geology of Minnesota: a centennial volume. Minnesota Geological Survey, University of Minnesota, Saint Paul, Minnesota.

Zhu, L., B. Fu, H. Zhu, C. Wang, L. Jiao, and J. Zhou. 2017. Trait choice profoundly affected the ecological conclusions drawn from functional diversity measures. Scientific Reports 7:3643.

